# SiteOut: an online tool to design binding site-free DNA sequences

**DOI:** 10.1101/029645

**Authors:** Javier Estrada, Teresa Ruiz-Herrero, Clarissa Scholes, Zeba Wunderlich, Angela H. DePace

**Affiliations:** Department of Systems Biology, Harvard Medical School, Boston, MA, USA.; School of Engineering and Applied Sciences, Harvard University, Cambridge, MA, USA.; Current address: Department of Developmental and Cell Biology, Center for Complex Biological Systems, University of California, Irvine, CA, USA.

## Abstract

DNA-binding proteins control many fundamental biological processes such as transcription, recombination and replication. A major goal is to decipher the role that DNA sequence plays in orchestrating the binding and activity of such regulatory proteins. To address this goal, it is useful to rationally design DNA sequences with desired numbers, affinities and arrangements of protein binding sites. However, removing binding sites from DNA is computationally non-trivial since one risks creating new sites in the process of deleting or moving others. Here we present an online binding site removal tool, SiteOut, that enables users to design arbitrary DNA sequences that entirely lack binding sites for factors of interest. SiteOut can also be used to delete sites from a specific sequence, or to introduce site-free spacers between functional sequences without creating new sites at the junctions. In combination with commercial DNA synthesis services, SiteOut provides a powerful and flexible platform for synthetic projects that interrogate regulatory DNA. Here we describe the algorithm and illustrate the ways in which SiteOut can be used; it is publicly available at https://depace.med.harvard.edu/siteout/.

## Introduction

Many essential biological processes depend on sequence-specific DNA binding proteins. It can be useful to test the functional role of individual binding sites or regulatory elements in the context of larger DNA sequences, but it can be difficult to design such perturbations. For example, during transcription, gene expression is controlled by transcription factors (TFs) that bind regulatory sequences such as enhancers and promoters (1). The role of individual TF binding sites in endogenous regulatory sequence, as well as their spacing and organization, is not well understood (2). A natural experiment to test the function of these DNA sequence features is to alter the number and arrangement of TF binding sites. Since transcription factor binding sites in animals are short and degenerate (the same protein may bind to multiple motifs; (1), they are easily created and therefore it is challenging to add, delete or move sites in a sequence without inadvertently affecting others. TF binding sites in isolation, e.g. in “neutral” sequence that lacks other binding sites have been tested using spacer sequences from orthogonal sources (e.g. bacterial and phage DNA used in *Drosophila*; (3, 4, 5) or synthetic random sequences (6, 7). However, it is virtually impossible to find orthologous or random sequences that entirely lack binding sites for proteins of interest (Figure 1). In specific instances the spacer sequence used can be shown to have no effect on expression levels (8). However, to rationally design and build complex sequences, it is useful to be able to control for binding site content directly. Manipulating short degenerate sequences is a general problem, common to studying other DNA binding proteins as well.

**Figure 1:**
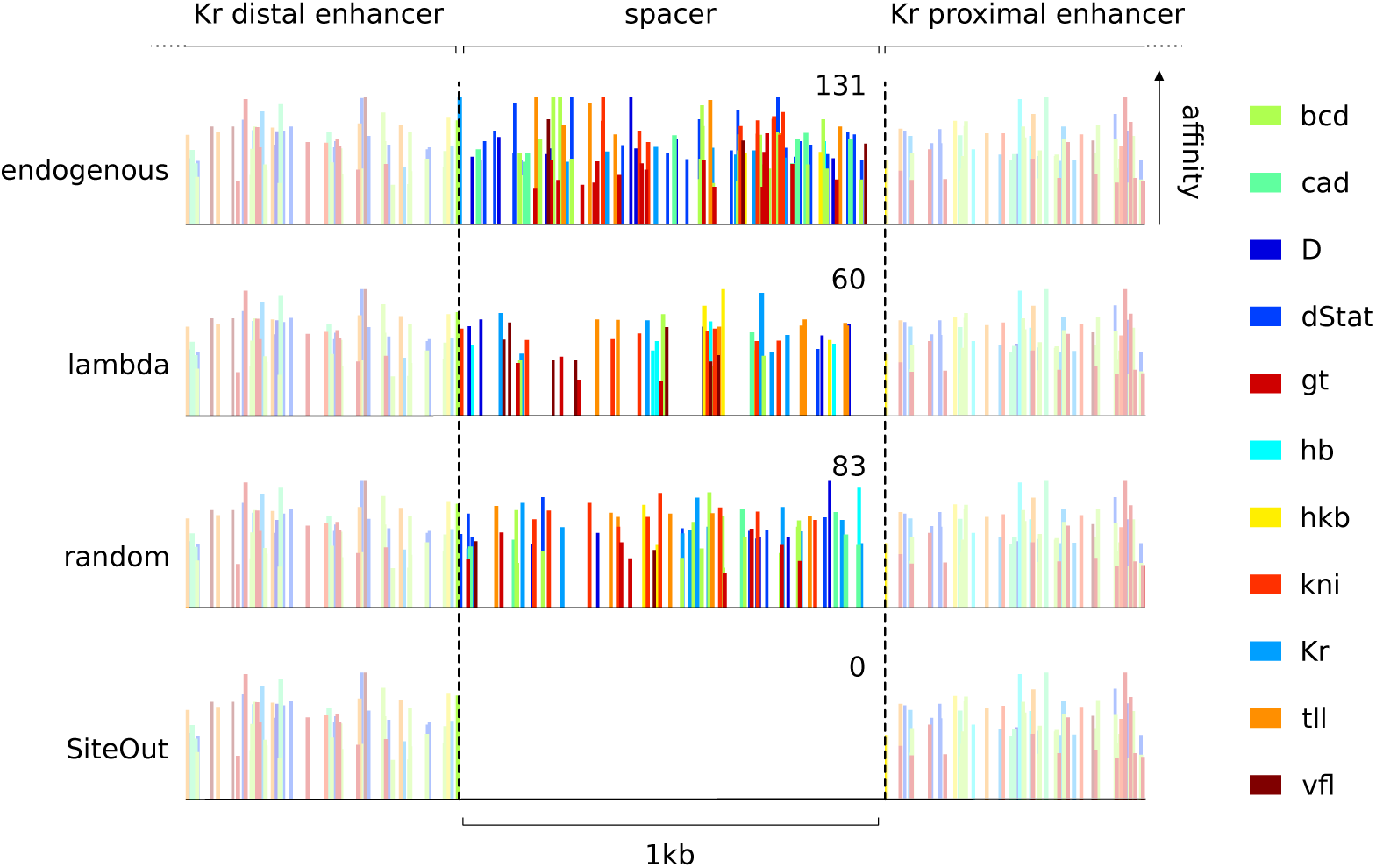
SiteOut efficiently removes binding sites from a spacer between annotated enhancers. Predicted binding sites for 11 transcription factors regulating expression of the gene *Krüppel* (Kr) in *Drosophila melanogaster* plotted across a 2kb region of its regulatory region; bar heights represent binding affinities. In the endogenous sequence between the two marked enhancers (labeled ‘spacer’) there are 131 binding sites for the transcription factors of interest. Sequences from orthogonal sources such as phage lambda have been used as “non-functional” spacers in many studies, but lambda DNA still contains many transcription factor binding sites. Randomly generated sequence with the same GC content as *D*.*melanogaster* intergenic sequence also contains a large number of binding sites. SiteOut creates a synthetic spacer that contains no binding sites, while keeping the flanking enhancers intact and the GC content constant.

Because it is now possible to synthesize arbitrary sequences at relatively low cost, we can easily make rationally designed DNA sequences to test specific hypotheses; the challenge is in designing sequence that contains few or no binding sites for proteins of interest. We have developed a tool called SiteOut to address this problem. SiteOut uses a Monte Carlo algorithm to iteratively remove all sites for proteins of interest from arbitrary sequence without generating new ones in the process. We provide examples to illustrate how SiteOut can be used to design synthetic enhancers and whole gene loci, and discuss the value of synthetic experiments for studying processes regulated by DNA-binding proteins more generally.

## Materials and methods

SiteOut begins with an initial sequence as defined by the user: a random sequence generated by SiteOut (*Random Sequence*), a sequence given by the user (*Refine a sequence*) or a combination of functional sequences and random spacers (Spacer Designer) (Figure 2A,B). The user specifies the GC content of the desired output sequence (GC content influences nucleosome positioning (9)) and the sites to be removed in the form of either specific sequences or frequency matrices, which are converted to Position Weight Matrices by the program (PWMs; (10)). In the latter case, PATSER (**stormo.wustl.edu**) is used to identify functional sites using a threshold p-value given by the user. SiteOut then removes binding sites iteratively using a Monte Carlo algorithm. This algorithm proves to be fast and reliable compared to other methods: a direct and deterministic search towards sequences that always lower the number of binding sites ends up stuck in local minima where not all sites are removed, while a genetic algorithm proves to be much slower (it has to work with multiple sequences) and ends up creating highly repetitive sequences. In each iteration, binding sites are identified and a random nucleotide in each site is mutated according to the nucleotide probability distribution given by the user-specified GC content (Figure 2C). An acceptance probability, *P*_a_, is computed to decide whether the mutated sequence is passed to the next iteration step. Mutations that decrease the total number of binding sites in the new sequence have a higher acceptance probability (eq.1: *N*_old_ and *N*_new_ represent the number of binding sites in the old and new sequences respectively). The algorithm stops once no binding sites are found, or when the sequence converges to a number of binding sites below which no fewer can be achieved within 100 iterations, which proves to be enough to avoid local minima. When the *Spacer Designer* option is used, the algorithm searches for sites that are within or overlap with the spacers and removes them by mutating only nucleotides from the spacer regions, keeping the functional sequences intact.

**Figure 2:**
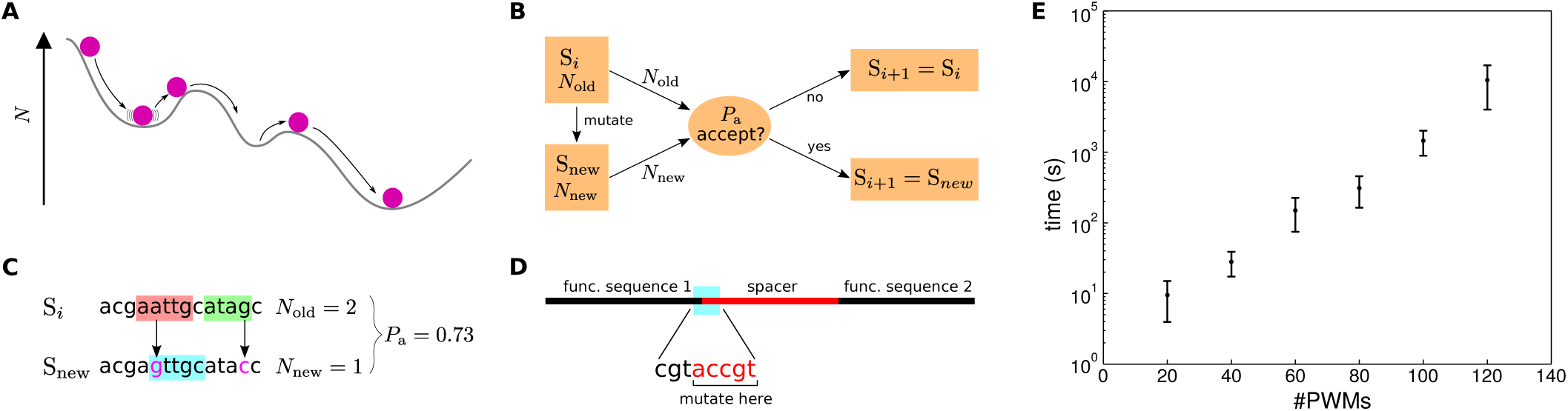
Overview of the SiteOut algorithm. (A) Schematic of the Monte Carlo algorithm. The non-deterministic nature of the process that drives the search towards sequences with fewer sites allows greater exploration of sequence space in order to overcome local minima in the number of binding sites (*N*). (B) Flowchart of the Monte Carlo algorithm highlighting the sequence acceptance/rejection process. Sm stands for a sequence in step *m* (*m = i, i* + 1,…), *N*_old_ and *N*_new_ for the number of binding sites in the original and mutated sequence, respectively, and *P*_a_ for the acceptance probability. (C) An example of binding site identification and deletion. Initially, two sites are identified (red and green), and are removed by mutating two random nucleotides (pink). This creates a new site (blue), but reduces the total number of sites in the sequence, thus giving an acceptance probability (*P*_a_) of 0.73. (D) Removing sites at the junction between a functional sequence and spacer by mutating only nucleotides from the spacer. (E) Performance plot for SiteOut running in Harvard Medical School’s cluster. Error bars come from different jobs being run in different nodes. Design of 300 bp random sequences, P value of 0.003. For 140 PWMs the 12 hour wall time is always reached.

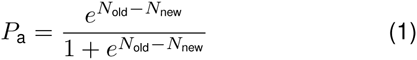

There are multiple trade-offs to consider when deploying SiteOut. First, SiteOut does not converge to a unique solution. For the same parameters, the algorithm will give a different solution every time. Therefore we advise that users perform multiple runs and choose the output that is most suitable for them. Second, the larger the number of binding sites to remove, the smaller the possible sequence space of the solution and the longer the run time (Figure 3E). Thus users need to carefully consider the length of the input sequence, how many motifs must be removed, and the threshold for binding site recognition (lowering the threshold will create more sites). Finally, if the output sequence is highly constrained, the algorithm is likely to generate repetitive sequences, which are challenging to synthesize and clone. This is not a limitation of SiteOut, but a byproduct of how easily binding sites are created and the small set of solutions when many constraints are applied. Working with these trade-offs, we have successfully used SiteOut to remove sites for 140 different yeast TFs and obtained a synthesizable binding site-free 1 kb sequence.

**Figure 3:**
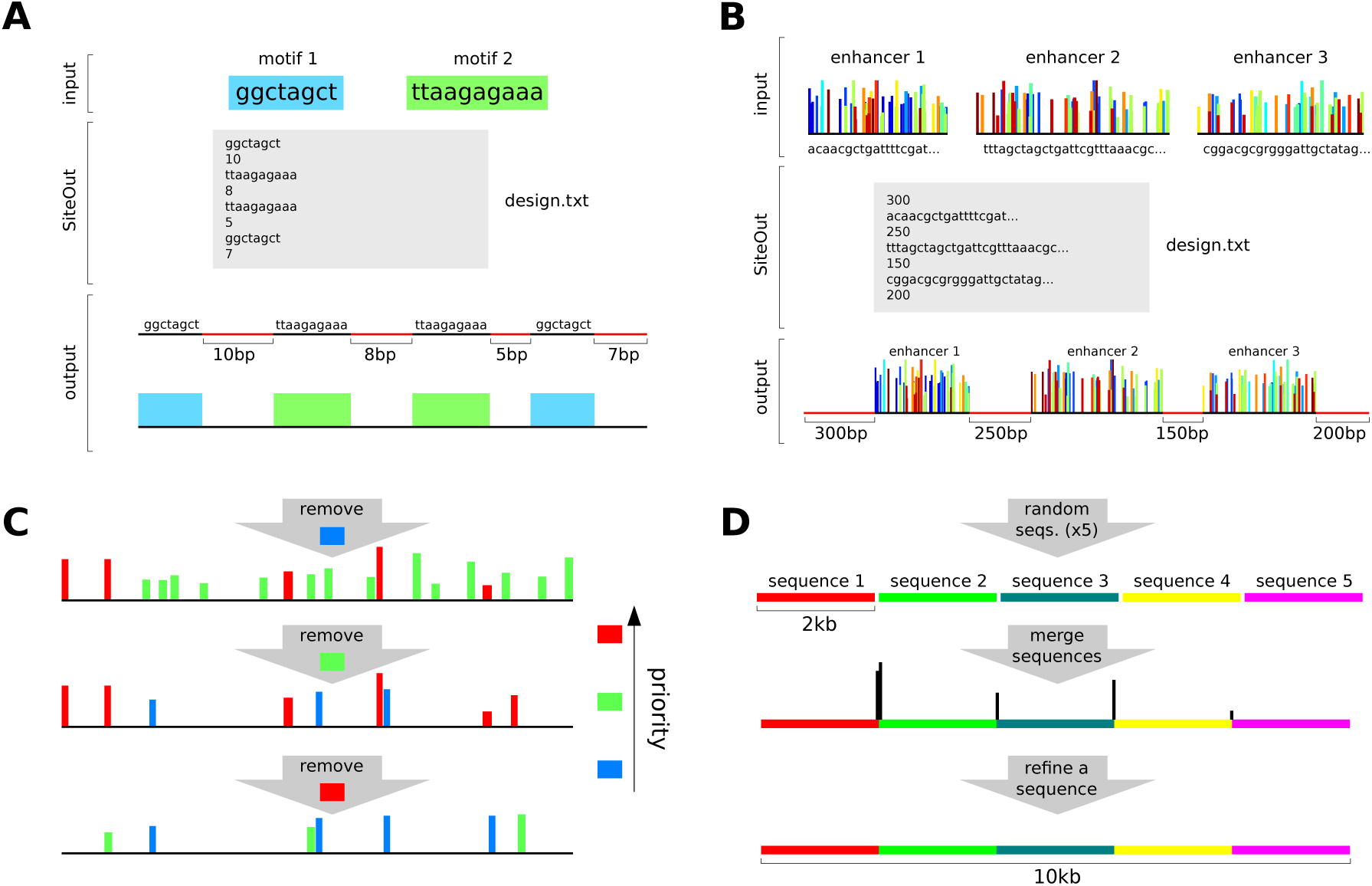
Examples of ways in which SiteOut can be applied. (A) Designing a synthetic enhancer. Binding motifs 1 and 2 are alternated forming a specific pattern to create a synthetic enhancer (bottom). The design.txt file shown in grey is the input given to SiteOut in this case. (B) Designing a synthetic gene locus. Three enhancers (top) are linked by 250 and 150 bp binding site-free sequences, and the whole construct is delimited by 300 and 200 bp site-free sequences. The design.txt file shown with grey background is the input given to SiteOut in the Spacer Designer option. (C) Removing binding sites in a hierarchical order if it is necessary to prioritize removal of particular sites. In this example we want to remove all red sites, most of the orange and as many as possible of the yellow. To do so, run SiteOut removing each binding site type one at a time, in reverse order of priority. In each step, the removal process may create new sites of a different type, but the most relevant ones are deleted in the subsequent steps. (D) Generating large binding site-free sequences by merging smaller ones. A 10 kb sequence (bottom) can be generated by merging 2 kb pieces made in parallel (top) and refining the resulting sequence (middle) to remove binding sites (black vertical bars) created at the junctions.

To allow users to add additional features to the algorithm’s screening process, we provide the source code on the website, so it can be downloaded and modified at will. For instance, the acceptance probability (*P*_a_) could be altered to include terms that take into account not only the number of binding sites in the sequence, but also their affinities or the number/length of repeats in the sequence.

## Tool description and examples

SiteOut presents the user with three main functionalities that confer enough flexibility to design a variety of DNA sequences (see the User Manual on the website):

1. **Design from scratch**: make a random binding site-free sequence of arbitrary length.
2. **Refine a sequence**: provide a sequence and remove specified binding sites.
3. **Design spacers in between functional sequences**: add spacers between sequences without creating binding sites at the junctions between the spacers and flanking functional sequences. Given a set of functional sequences that will remain untouched, SiteOut links them together using binding site-free spacers of the desired lengths (these can be from a few nucleotides to many kilobases long).

Here we illustrate how to use the tool with four examples from transcription:

- **Designing a synthetic enhancer:** Testing if a set of experimentally-identified or computationally-predicted binding sites is sufficient to drive a specific gene expression pattern is challenging. The *Spacer Designer* can be used to do this by designing site-free sequences between known binding sites (Figure 3A).
- **Designing a synthetic gene locus:** The activity of regulatory elements in a gene locus can be assayed in isolation from their endogenous sequence surroundings by using the *Spacer Designer* to generate site-free sequences between them. The algorithm avoids generating any of the specified binding sites at the junction between functional sequence and spacer (Figure 3B).
- **Removing binding sites in a hierarchical order:** There may be situations where the number of binding sites to be removed is so high that no zero-binding site solution exists. In this case one may want to prioritize the removal of some binding sites over others. This can be done by running SiteOut several times, removing one type of binding site at a time, in order of increasing importance (Figure 3C).
- **Generating large sequences by merging smaller ones:** When designing very long binding site-free sequences it is convenient to parallelize the job to save time and avoid hitting the 12-hour maximum run time set on the Harvard Medical School computing cluster. For example, if designing a 10 kb sequence it is much faster to submit five jobs using the *Random Sequence* option, each designing a different 2 kb piece. These five sequences can be connected together and the resulting sequence refined using the *Refine a Sequence* option, which will remove any sites that formed at the junctions between them (Figure 3D).

## Discussion

Synthetic approaches to dissecting the activity of DNA-binding proteins are more tractable than ever, given the diminishing price of synthesizing DNA sequence (11). This opens up wide-ranging opportunities to test specific hypotheses about how the identity, number and arrangement of protein binding sites in regulatory DNA enable precise control over biological processes. For example, models of gene regulation rely on computationally-predicted transcription factor binding sites, but these sites may not faithfully reflect in vivo protein binding (12, 13, 14) and binding sites outside predicted enhancers can be essential for transcriptional regulation (15). It is therefore valuable to explicitly test the importance of binding site content, as opposed to other sequence features of DNA such as its mechanical properties, shape, chromatin structure or the spacing between regulatory elements. As we better understand how DNA sequence maps to in vivo binding site occupancy and protein function, additional features can be controlled for in later versions of Site-Out.

While we have focused on transcription to illustrate the use of SiteOut, the ability to design carefully controlled binding site-free sequence will be useful for investigating many other process controlled by sequence-specific DNA-binding proteins. For instance, DNA replication in bacteria is controlled by a host of factors that recognize specific sites in the origin and regulate DnaA recruitment to binding that differ in affinity, spacing and orientation across species. It is unclear how differences in the structure of the replication origin map onto species-specific features of DnaA oligomerization and its control by other regulatory proteins (16), but this question is ripe for the types of synthetic experiments discussed here.

## Acknowledgments

We thank the members of the DePace Lab for their thoughtful comments and ideas during SiteOut development, with a special acknowledgment to Ben Vincent, who came up with such an eloquent name. We also acknowledge Daniel Navarro, at the Mass. Eye and Ear Infirmary, for his help on setting up the web interface.

